# Do aye-ayes echolocate? Studying convergent genomic evolution in a primate auditory specialist

**DOI:** 10.1101/048165

**Authors:** Richard J. Bankoff, Michael Jerjos, Baily Hohman, M Elise Lauterbur, Logan Kistler, George H. Perry

## Abstract

Several taxonomically distinct mammalian groups – certain microbats and cetaceans (e.g. dolphins) – share both morphological adaptations related to echolocation behavior and strong signatures of convergent evolution at the amino acid level across seven genes related to auditory processing. Aye-ayes (*Daubentonia madagascariensis*) are nocturnal lemurs with a derived auditory processing system. Aye-ayes tap rapidly along the surfaces of dead trees, listening to reverberations to identify the mines of wood-boring insect larvae; this behavior has been hypothesized to functionally mimic echolocation. Here we investigated whether there are signals of genomic convergence between aye-ayes and known mammalian echolocators. We developed a computational pipeline (BEAT: Basic Exon Assembly Tool) that produces consensus sequences for regions of interest from shotgun genomic sequencing data for non-model organisms without requiring *de novo* genome assembly. We reconstructed complete coding region sequences for the seven convergent echolocating bat-dolphin genes for aye-ayes and another lemur. Sequences were compared in a phylogenetic framework to those of bat and dolphin echolocators and appropriate non-echolocating outgroups. Our analysis reaffirms the existence of amino acid convergence at these loci among echolocating bats and dolphins; we also detected unexpected signals of convergence between echolocating bats and both mice and elephants. However, we observed no significant signal of amino acid convergence between aye-ayes and echolocating bats and dolphins; our results thus suggest that aye-aye tap-foraging auditory adaptations represent distinct evolutionary innovations. These results are also consistent with a developing consensus that convergent behavioral ecology is not necessarily a reliable guide to convergent molecular evolution.

## Introduction

The aye-aye (*Daubentonia madagascariensis*) is an Endangered nocturnal lemur that is the only surviving member of the family Daubentoniidae. Among other unique adaptations, aye-ayes possess an elongated and highly flexible middle finger with which they tap rapidly along the surface of trees in the search for the internal mines of wood-boring insect larvae (Erickson 1991; Sterling and Povinelli 1999; Schwitzer et al. 2013; Thompson et al. 2016). The resulting differential soundings of a tree’s variable interior structures are received by aye-ayes' large, alert, high-frequency attuned pinna (Coleman and Ross, 2004; Ramsier and Dominy, 2012) and processed by a relatively enlarged inferior colliculus in a brain that is also overall the largest relative to body size among all extant strepsirrhine primates (Smith and Jungers 1997; Kaufman et al. 2005). In addition to acoustic signals, tactile and olfactory cues are also hypothesized to play important roles in aye-aye foraging ecology (Erickson, 1991; Bush and Allman, 2004; Kaufman et al., 2005; Ramsier and Dominy, 2012). Once a suitable location on the deadwood is detected, the aye-aye uses its large, continuously-growing incisors to gnaw through the tree's exterior and extract the larvae within using the flexible middle finger (Erickson, 1991; Soligo, 2005).

The tap-foraging adaptations of the aye-aye have led to analogies to woodpeckers, as a result of their similar extractive, insectivorous dietary niche (e.g. (Sandwith, 1859; Cartmill, 1974). Woodpeckers investigate trees for potential food sources by systematically probing and excavating cavities within a copse for grubs before moving on to a new area (Lima, 1983). To test whether aye-ayes conform to expectations of systematic and/or random cavity investigation, Erickson (1991) conducted a series of behavioral experiments with four wild-caught, captive aye-ayes at the Duke Lemur Center. The aye-ayes were significantly more likely to excavate cavities relative to non-hollow areas of the block, regardless of whether the cavities contained larvae (Erickson, 1991), indicating that aye-ayes can consistently distinguish true cavities from false indicators even in the absence of visual or olfactory cues by using the acoustic signals generated in tap-foraging to identify potential larval mines.

The neurological and genetic pathways involved in aye-aye tap-foraging behavior are currently unknown. Recent molecular analyses of ecologically and morphologically convergent, but phylogenetically distinct, taxa have shown that – at least in some cases – similar genetic changes may underlie convergent biology (Stern 2013; Gallant et al. 2014). For example, *RNASE1* gene duplications and subsequent amino acid substitutions occurred independently in folivorous colobine primates and ruminating cows, likely facilitating the digestion of large amounts of bacterial RNA in lower pH conditions associated with foregut fermentation (Zhang et al. 2002; Zhang 2003; Schienman et al. 2006; Zhou et al. 2014). The availability of aye-aye genomic sequence data (Perry et al. 2012; Perry et al. 2013) facilitates cross-species analyses that may contribute to our understanding of the underlying biology.

Our present analysis is motivated by the observation of molecular convergence among echolocating bats (two phylogenetically divergent clades; suborders Yinpterochiroptera and Yangochiroptera) and between echolocating bats and the toothed whales (suborder Odontoceti). Bat and whale echolocators exhibit radical differences in their mechanisms of sound production, with the former relying on the standard mammalian laryngeal apparatus to emit vocalizations and the latter possessing a specialized nasal structure for sound production (Cranford et al. 1996; Au and Simmons 2007; Jakobsen et al. 2013). Despite using distinct organs for producing and propagating sounds, a striking pattern of convergent genomic evolution in seven genes involved in auditory functioning suggests similarities in at least some of the ways that the returning sounds are processed. Specifically, there is strong consensus evidence for seven auditory processing genes with convergent amino acid substitutions among the echolocating bats and dolphins (**Table 1**), to such a degree that phylogenetic reconstructions of the predicted protein sequences of these genes produce monophyletic clades of all echolocators to the exclusion on of their more closely related, non-echolocating sister taxa (Liu et al. 2010; Shen et al. 2012). While recent suggestions of an even wider, cross-genome level of convergence (Parker et al. 2013) have not been supported by subsequent analyses (Thomas and Hahn 2015; Zou and Zhang 2015), the evidence for convergence at the seven genes listed in **Table 1** is robust.

**Table 1.**
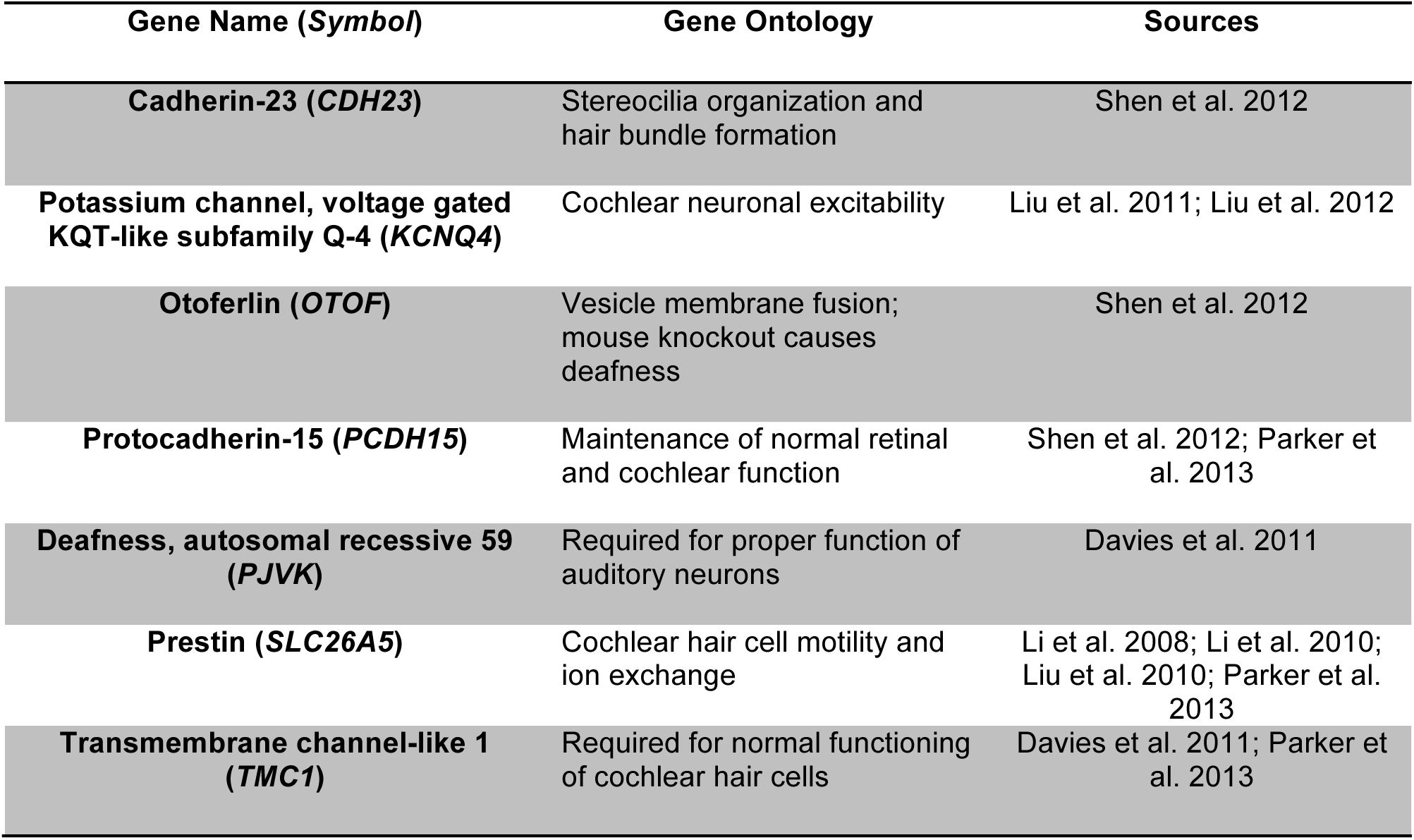
Genes previously shown to have significantly convergent encoded amino acid sequences among echolocating bats and dolphins

In this study, we tested whether the genomic signal of convergent adaptation detected between the echolocating bat and dolphin lineages was shared with aye-ayes. Given that these patterns of convergence appear to be organized around a common reliance on interpreting complex auditory signals to detect prey in different media rather than mechanisms of sound production, we hypothesized that the relatively poorly understood neurological and mechanical pathways that aye-ayes use to process high-frequency acoustic signals might have similar genomic underpinnings. To test this hypothesis, we reconstructed all seven loci implicated in previous bat dolphin comparisons in aye-ayes and the diademed sifaka (*Propithecus diadema*), a sister taxon, using both new and previously published genomic short read data (Perry et al. 2012; Perry et al. 2013) and the Basic Exon Assembly Tool (BEAT), a newly developed pipeline that links multiple existing bioinformatics tools to rapidly extract consensus sequences from loci of interest from shotgun sequence data. We then evaluated the level of amino acid convergence among these lemurs and 12 additional species including echolocating bats and dolphins and their non-echolocating sister taxa.

## Material and Methods

### Data Acquisition

For aye-aye sequences, sequence short reads from Perry et al. (2012) and Perry et al. (2013) were retrieved from the National Center for Biotechnology Information (NCBI) Sequence Read Archive (www.ncbi.nlm.nih.gov/Traces/sra, accession numbers SRA066444 & SRA043766.1) and queried as compressed FASTQ files, comprising 32 lanes of data (3,842,334,284 reads). Tissue samples from two *Propithecus diadema* individuals were provided by the Duke Lemur Center. Sifaka sequencing libraries were prepared using the method described in Meyer and Kircher (2010). Each individual was sequenced on one lane of the Illumina HiSeq 2500, for 150 cycles from each end (150x150 paired end sequencing; 547,279,426 reads in total for the two individuals). The diademed sifaka sequence read data generated for this study have been deposited in the Sequence Read Archive with the BioProject accession number PRJNA317769.

In addition to the aye-aye and sifaka data, our analysis included *CDH23, KCNQ4, OTOF, PCDH15, PJVK, SLC26A5* (*Prestin*), and *TMC1* (see **Table 1**) gene sequence data that were provided by Parker et al. (2013) for 12 species, including six bats (echolocators: *Pteronotus parnellii, Megaderma lyra, Rhinolophus ferrumequinum*, and *Myotis lucifugus*; non-echolocators: *Eidolon helvum* and *Pteropus vampyrus*), bottlenose dolphins (*Tursiops truncatus*) and their non-echolocating relative domestic cattle (*Bos taurus*), and three mammalian outgroups: house mouse (*Mus musculus*), African elephant (*Loxodonta africana*), and European rabbit (*Oryctolagus cuniculus*). *TMC1* was not available for African elephants, so analyses involving this species are based on the six remaining genes. Other species from the original Parker et al. (2013) alignments with less than 75% sequence length coverage for any gene were excluded from our analyses. The original Parker et al. (2013) alignments were missing *SLC26A5* for *Eidolon helvum*; however, we were able to reconstruct this gene using short read data from reads accessioned by Parker et al. (2013) (NCBI accession number SRA091593) as the input and a *Pteropus vampyrus* ortholog. Gene nomenclatural differences between Parker et al.'s (2013) alignments and the NCBI's nuccore database used for the assembly of *Daubentonia madagascariensis* and *Propithecus diadema* sequences were standardized using the UCSC table browser (https://genome.ucsc.edu/cgi-bin/hgTables). Where more than one isoform of a gene has been reported, the homologous human sequences that *Daubentonia madagascariensis* and *Propithecus diadema* short read sequences were mapped to with BEAT were chosen to match the isoforms used in the multi-species alignments from Parker et al. (2013).

### Data Generation

The Basic Exon Assembly Tool (BEAT, https://github.com/RBankoff/BEAT) is a set of perl shell scripts intended to quickly and reliably reconstruct a consensus sequence from a non-model organism for a genomic region that is orthologous to a reasonably-conserved reference sequence from the genome of a distantly-related species, using a set of raw Illumina paired-end short read data from the non-model species with sufficient depth of coverage to reliably call genotypes. The capability to easily and accurately create assemblies for non-model species such as the aye-aye is critical to maximize data usability for endangered or otherwise difficult-to-study species, particularly for research groups not explicitly focused on bioinformatic analyses.

BEAT acts as a pipeline to coordinate the parallelization of multiple freely available programs (released under the GNU GPL3 License) in a UNIX/LINUX environment, as described below. BEAT does *not* introduce any novel techniques for assaying genomic information for quality or analyzing the data it generates; its purpose is to facilitate the efficient extraction of selected regions of interest from a large but otherwise unassembled set of raw reads in a single end-user step using parallelization of computationally intensive processes. BEAT can be run through the GNU parallel shell tool (Tange 2011) on a workstation or can take advantage of distributed cluster computing environments such as the TORQUE Process Manager to map multiple sets of paired-end short read sequence data from different flowcells and individuals sequenced on an Illumina platform to a high-quality reference genome in parallel. By linking directly to the Entrez database with the free EntrezDirect tool from NCBI (Kans 2011), BEAT can take user requests for specific loci and obtain reference coordinate and sequence information for orthologous sequences from a specified reference genome.

BEAT maps reads to either a reference genome (e.g. GRCh38) or other reasonably unique sequence(s) of interest by running the BWA-MEM algorithm (Li 2013) over each set of provided paired-end reads in parallel (see **Figure 1**). For example, a BEAT job that uses 20 paired-end short read files will be submitted to run as 10 simultaneous jobs. For larger short read datasets, BEAT will subdivide further, breaking them into small files (default 1GB) that can be further processed in parallel. The raw .bam-formatted mapped output files are quality filtered to exclude unmapped reads with the samtools view ‐F4 flag (Li et al. 2009) and passed through samtools rmdup to remove PCR duplicate reads that might otherwise bias consensus calls, before being merged into a master .bam alignment file. Per-nucleotide genotype estimates are then generated using the samtools mpileup utility, which calls the read-specific genotype and read quality from the .bam alignment file into a position-wise pileup output file. This master pileup file can subsequently be queried for the corresponding sequence call at any position along the length of its reference using the BEAT_consensus script. A novel component of BEAT, this script takes advantage of the read-by-read genotypes mapped to the ortholog in the mpileup output file to generate a consensus sequence based on the majority genotype across all reads mapping to a particular range of coordinates along the orthologs. The creation of consensus sequences is accompanied by the generation of a summary statistics file containing per sub-feature (e.g. exon, promoter region, or other segment of interest) coverage, using the sum of position-wise read depth / length of sub-target. As we show below, consensus sequence output data produced by BEAT for a given region is more complete than that contained within a corresponding genome assembly from the same sequencing data, while requiring significantly lower computational investments.

**Figure 1.**
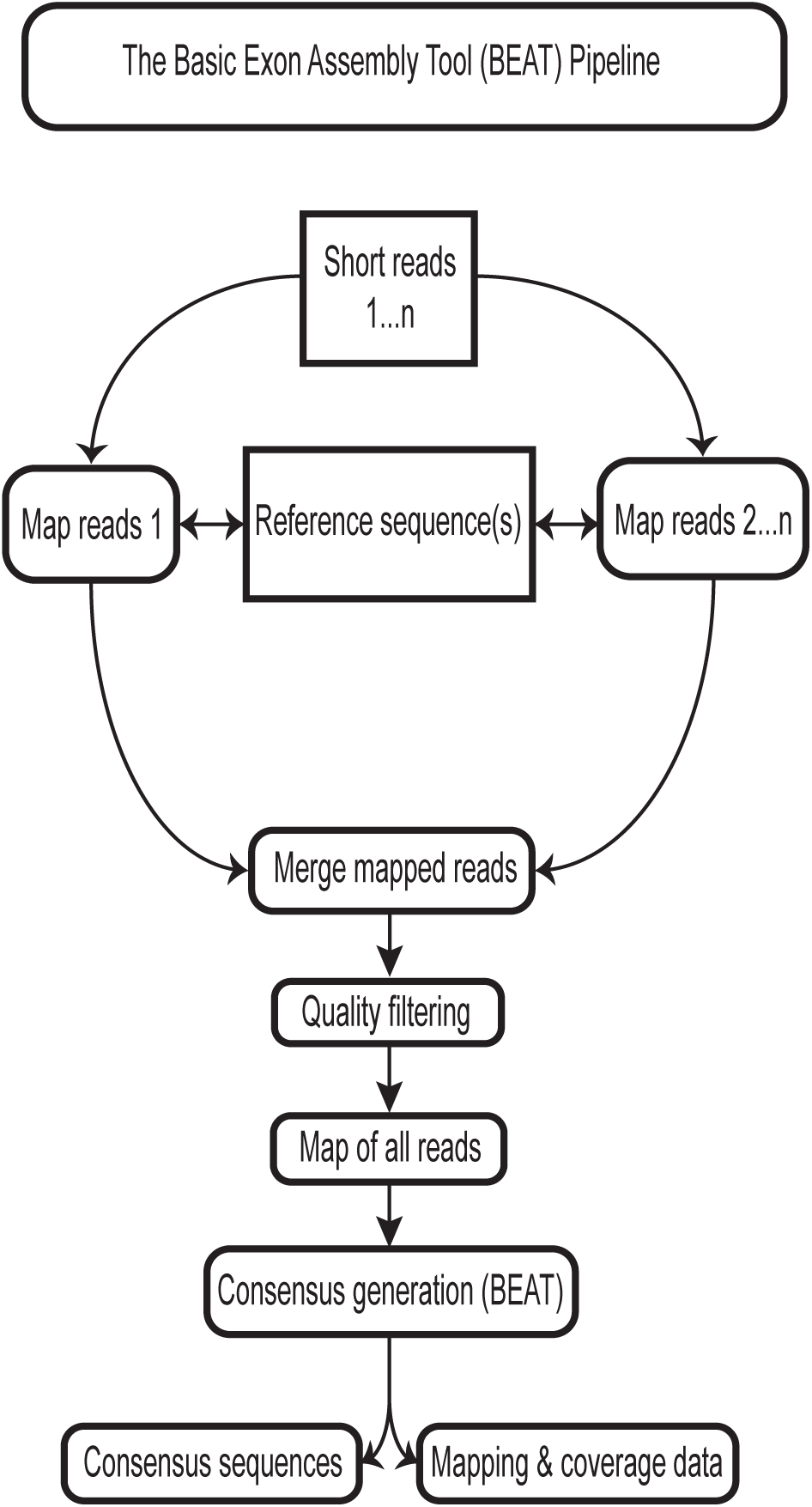
The basic workflow used by BEAT when assembling a single query sequence from a short read dataset. BEAT maps all short reads provided to a reference genome in parallel with BWA-MEM, then merges the mapped files to produce a map of the entire reference sequence that can be searched repeatedly. The resulting mapped files are passed through the BEAT consensus generator to produce consensus sequences for loci of interest.

The ideal parameters for running the BEAT pipeline include 16gb RAM and one computing core per set of ~20gb combined compressed short read data files in FASTQ format. Lower-performance systems should still be able to run BEAT, but job runtime may differ significantly from those reported below, as parallelization is the key to BEAT's efficiency.

### Assessment of BEAT performance

To test the targeted region assembly performance of BEAT compared to that of a *de novo* complete genome assembly generated using the same underlying sequence data, we assessed the percent coverage of consensus sequences generated by BEAT to that achieved by locating matching sequences using the blastn utility in BLAST+. We mapped *Daubentonia madagascariensis* short read data to the human orthologs of each of the seven genes of interest using BEAT and compared the percent coverage to that achieved by locating matching sequences using BLAST from the aye-aye genome assembly scaffolds published in Perry et al. (2012). While the consensus sequences generated by BEAT for the convergence analysis presented in the Results of this paper used additional short read data from Perry et al. (2013), for this assessment, consensus sequences were regenerated solely from the read data used to create the original scaffolds published in Perry et al. (2012) to achieve full BEAT-scaffold comparability.

For the BEAT-generated data for this test, aye-aye short reads from Perry et al. (2012) were processed using the human orthologs from the Parker et al. (2013) alignments as BLAST baits. Short reads that matched these orthologs were then mapped to the orthologous sequences using BEAT, and a consensus sequence was generated in BEAT_consensus for any nucleotides with a minimum of two uniquely mapping reads that passed the minimum mapping quality threshold. For sequences from the Perry et al. (2012) scaffolds, the same human orthologs for all seven genes were queried using blastn against the scaffolds with a minimum alignment score of 50, identity cutoff set at 80%, and a word length of 11. Sequences that BLAST matched to the orthologs were extracted from the scaffolds and aligned to the ortholog in Geneious. Following BLAST hit extraction, BEAT mapping, and alignment to the ortholog in Geneious, per-gene percent coverage of scaffold‐ and BEAT-derived sequences was calculated as the percentage of nucleotides in the ortholog with a corresponding aligned (though not necessarily identical, given expected human and aye-aye sequence divergence) nucleotide from the test datasets. We then compared the percent coverage results for the seven BEAT-generated sequences to those for the scaffold-generated sequences. We found that BEAT retrieves more sequence data on average than using blastn to query the *de novo* assembly genomic scaffolds from Perry et al. (2012), with an overall mean 99.75% coverage for consensus sequences generated by BEAT versus 91.13% mean coverage for querying the genome assembly scaffolds with BLAST (Table 2).

**Table 2.** Comparison of *de novo* genome assembly and BEAT methods for reconstruction of aye-aye gene coding regions using the same shotgun sequence read data for both methods.

| **Gene** | **<u>Human</u>**<br>**<u>reference</u>**<br>**<u>coding region</u>**<br>**<u>size (bp)</u>** | **<u>Aye-aye *de novo* assembly</u><sup>1</sup>** |  | **<u>BEAT with aye-aye sequence</u>**<br>**<u>reads</u><sup>2</sup>** |  |
| --- | --- | --- | --- | --- | --- |
|  |  | bp recovered | % coverage | bp recovered | % coverage |
| *CDH23* | 8979 | 8122 | 90.46 | 8909 | 99.18 |
| *KCNQ4* | 1713 | 1192 | 69.59 | 1713 | 100 |
| *OTOF* | 5634 | 4587 | 81.42 | 5632 | 99.96 |
| *PCDH15* | 4195 | 4120 | 98.21 | 4194 | 99.98 |
| *PJVK* | 849 | 843 | 99.29 | 842 | 99.18 |
| *SLC26A5* | 2040 | 2040 | 100 | 2040 | 100 |
| *TMC1* | 2124 | 2102 | 98.96 | 2124 | 100 |

<sup>1</sup> Analysis based on BLAST reconstructions of scaffolds from aye-aye *de novo* assembly published in Perry et al. (2012).
<sup>2</sup> Analysis based on BEAT reconstructions from the same shotgun sequencing read data used in Perry et al. (2012), a subset of the sequence read data used in the full analysis of this paper.

### Data processing and analysis

For the evolutionary analyses performed in this study, consensus sequences for seven echolocation convergence candidate genes for both *Propithecus diadema* and *Daubentonia madagascariensis* were generated with the BEAT pipeline, using human RefSeq mRNA sequences (assembly hg19/GRCh38) as the input reference orthologous sequences (**Table 1**). Consensus calls for each nucleotide were calculated at each site with ≥2x unique mapped read coverage. Consensus sequences for each gene were generated on an exon-by-exon basis, aligned to the canonical mRNA transcript version of the human ortholog using Geneious version 8.0.5 (http://www.geneious.com, Kearse et al. 2012), and assembled into final consensus sequences for the coding sequence of each gene and species. These final consensus sequences were aligned with the 14 species nucleotide alignments from Parker et al. (2013) in Geneious using the Geneious alignment algorithm, with a high gap open penalty of 25 but a low gap extension score of 2 to discourage fragmentation while allowing for indels. These alignments were manually assessed for quality, including removal of any codon frame-altering indels relative to the human reference sequence. Curated nucleotide alignments for each gene were translated into predicted amino acid sequences in Geneious. The consensus amino acid sequence for each gene was determined with a strict 50% cutoff (if the most common amino acid was observed in fewer than 50% of the species in the alignment, then no consensus was recorded for that position). All final multi-species amino acid and DNA alignments used in this study have been deposited in the Open Access Scholarsphere digital repository at https://scholarsphere.psu.edu/collections/rv042t11v.

To evaluate the level of amino acid convergence across the seven candidate genes between *Daubentonia* and the echolocating bat and dolphin lineages, we devised a straightforward method to correct for the effects of phylogeny on patterns of convergence, illustrated with an example in **Figure 2**. First, consider a simplified set of three species from the alignment: for example, *Daubentonia, Propithecus*, & *Tursiops*, where *Daubentonia* and *Propithecus* are more closely related to each other than either is to *Tursiops*. In the absence of convergent evolution between dolphins (*Tursiops*) and one of the two lemur species (*Daubentonia, Propithecus*), *Tursiops* is expected to share an approximately equal number of identical, aligned amino acids at each gene with *Daubentonia* but not *Propithecus* as it is with *Propithecus* but not *Daubentonia*. Only sites with shared amino acids between *Daubentonia* and *Tursiops* that also differ from both the *Propithecus* sequence and the multi-species consensus sequence are considered convergent. In other words, we count the number of sites at which *Tursiops* shares an amino acid with *Daubentonia* but not with *Propithecus* or the multi-species consensus sequence, relative to the total number of amino acids in the alignment. We then compute a ratio by comparing this result to the count of the number of amino acid shared by *Tursiops* and *Propithecus* but not with *Daubentonia* or the consensus (again, relative to the total number of aligned amino acids). This ratio can be computed for any species pair-outgroup combination, can be computed for an individual gene or summed across multiple genes, and can be statistically evaluated with a Fisher’s Exact Test. For the purposes of this analysis, we summed all convergent amino acids and divided by the sum of all amino acids across all seven genes for each species pair-outgroup combination.

**Figure 2.**
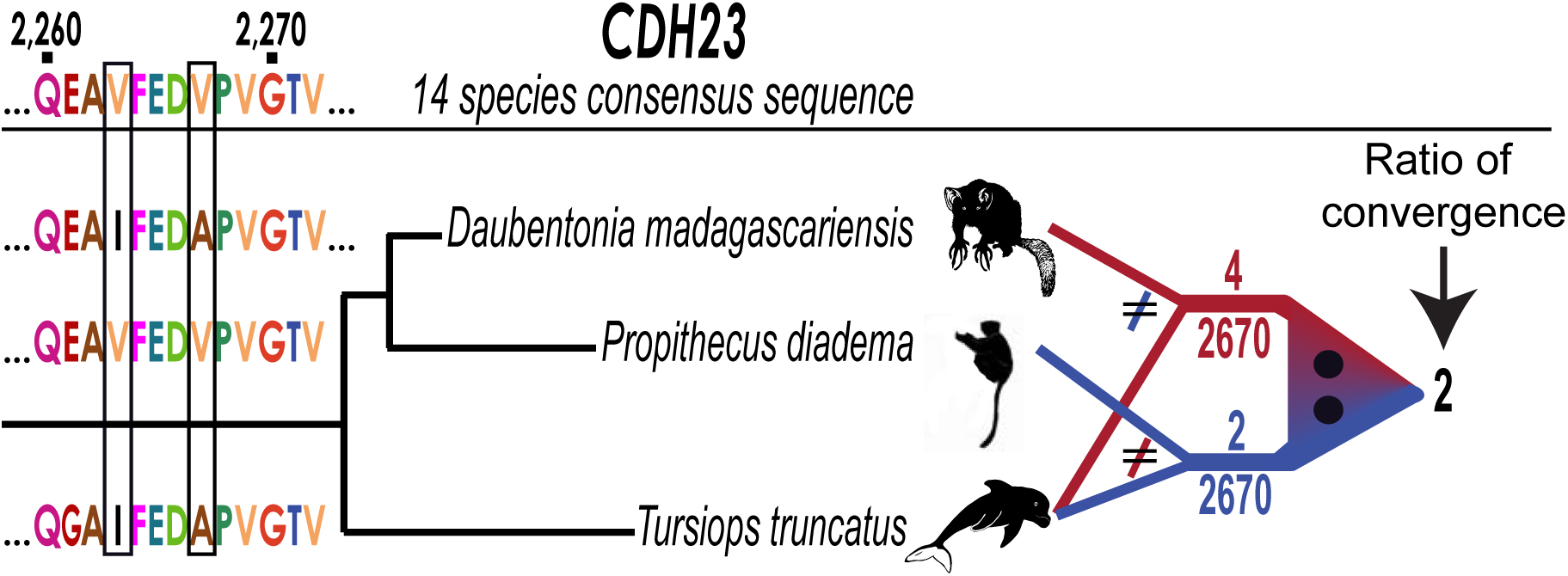
Illustration of the pairwise convergent evolution analysis used in this study. The example shown is for a section of the *CDH23* gene, here comparing two lemurs, *Daubentonia* and *Propithecus*, to the echolocating dolphin *Tursiops*. For each lemur, we computed the number of positions out of the total aligned positions (2,670 for this gene) for which that species shared an amino acid with *Tursiops*, but for which that amino acid was different than that of the other lemur species and the consensus sequence determined from the multiple-species alignment. A section of *CDH23* is shown that contains two *Daubentonia* – *Tursiops* convergent amino acids. For the whole gene, the ratio of convergent amino acids for *Daubentonia* and *Tursiops* (4) and *Propithecus* and *Tursiops* (2) = 2, providing a magnitude and directionality of relative convergence between aye-aye and dolphin, with a phylogenic correction based on the sister lemur species. These ratio values are depicted in **Figure 3** for all tested comparisons, summed across the seven genes analyzed in this study. The difference in the number of convergent amino acids is evaluated with a Fisher’s exact test; for the illustrated *CDH23* comparison, *P* = 0.6873, prior to Bonferroni multiple test correction.

To test the hypothesis of genomic convergence between aye-ayes and the true echolocators with this method, we followed the above procedure separately for each echolocating species, i.e., we compared the ratio of the proportion of *Daubentonia* and echolocator convergent amino acids (to the exclusion of *Propithecus* and the consensus) to the proportion of *Propithecus* and echolocator convergent amino acids (to the exclusion of *Daubentonia* and the consensus), separately for each echolocating species in the multi-species alignment. Similarly, we compared bottlenose dolphins (*Tursiops truncatus*) and a number of echolocating bat species to one another using domestic cattle (*Bos taurus*) and non-echolocating bats, respectively, as the non-echolocating sister species in the analysis. For each pairwise+outgroup comparison, only sites without gaps or missing data in all three species’ sequence (and a called consensus amino acid) were counted toward the total number of amino acids used in calculating the ratio. To correct for the number of tests performed, we applied a Bonferroni correction by multiplying the individual Fisher’s Exact Test p-values by the number of tests.

## Results

Echolocation has likely evolved twice independently in bat species (Jones and Teeling 2006; Jones and Holderied 2007) and at least once in toothed whales (Geisler et al. 2014). Previous studies have shown that echolocating bats and dolphins share significantly more amino acids with each other at seven genes implicated in auditory processing than they do with phylogenetically closer non-echolocating taxa (Liu et al. 2011; Shen et al. 2012). These previous observations are strongly supported by the results from our echolocator vs. non-echolocating sister group pair-wise comparisons for these seven genes (Figure 3). For example, dolphins (*Tursiops truncatus*) shared 65 convergent amino acids (out of 7,734 total aligned positions across the seven studied genes) with the echolocating bat *Rhinolophus ferrumequinam* to the exclusion of cattle (*Bos Taurus*, more closely related to dolphins), whereas *Rhinolophus* and *Bos* shared only 7 convergent amino acids to the exclusion of *Tursiops*, for a convergence ratio = 9.3, which is significantly greater than expected by chance (Fisher’s exact test, *P* = 6.402E-13; Bonferroni-corrected *P* = 2.816E-11). Similarly, the echolocating bat *Rhinolophus ferrumequinam* and *Tursiops* share 65 out of 7,655 comparable sites, while *Tursiops* and *Eidolon helvum*, a non-echolocating bat, share only 4 (convergence ratio = 16.3; *P* = 2.844E-15; Bonferroni-corrected *P* = 1.251E-13). Significant convergence was observed between all other echolocator-echolocator comparisons, but not between dolphins and non-echolocating bats (**Figure 3**; **Supplemental Table 1**). On a gene-by-gene basis, *CDH23, OTOF, PCDH15*, and *SLC26A5 (Prestin)* display the strongest, most consistent signals of convergence in the echolocator-echolocator comparisons (see **Supplemental Table 2** for gene-by-gene comparison results).

**Figure 3.**
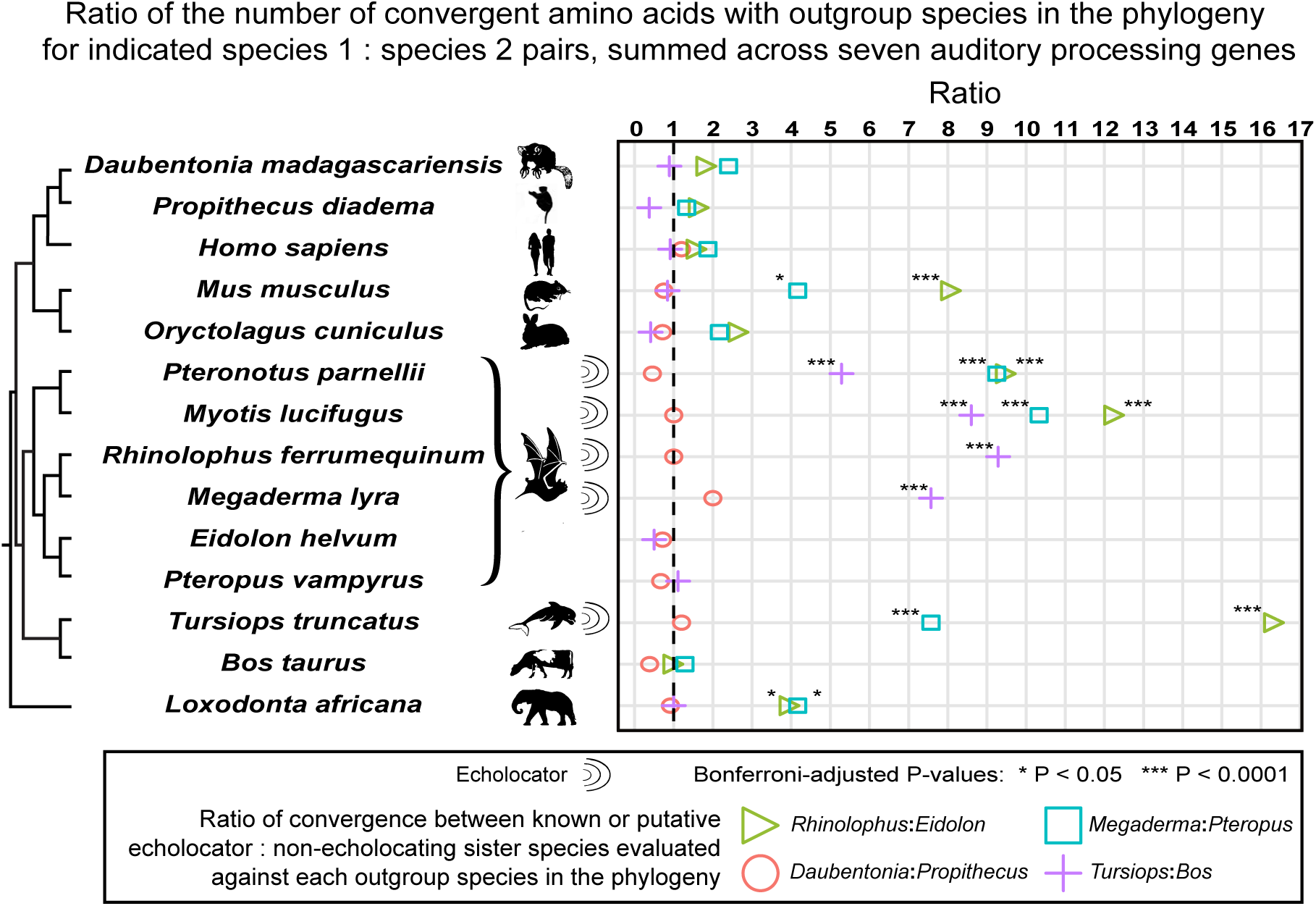
Observed seven gene convergence ratios for three pairs of known echolocator:non-echolocator sister taxa and for *Daubentonia:Propithecus*, each compared to other echolocating and non-echolocating mammalian taxa. Colored symbols represent the ratio between the number of convergent amino acids shared between the first species in the indicated pair and the species on the corresponding row of the phylogeny (the “test species”) relative to that for the second species in the indicated pair, after results were summed over the seven auditory genes considered in this analysis. The dashed line indicates a ratio of 1, or no difference in the observed level of convergence with the outgroup test species between the two sister species. The indicated significance values have been corrected for multiple tests using the Bonferroni method, within each set of pairwise comparisons across the phylogeny.

In contrast to the significant results for the known echolocator vs. known echolocator lineages, we did not observe a signature of convergent amino acid evolution between *Daubentonia* and the echolocating bats or dolphins relative to *Propithecus* (**Figure 3; Supplemental Table 1**). For example, in sum across the seven genes, *Daubentonia* and *Rhinolophus* shared 10 convergent amino acids (of 8,235 positions) to the exclusion of *Propithecus*, while *Propithecus* and *Rhinolophus* also shared 10 convergent amino acids to the exclusion of *Daubentonia* (convergence ratio = 1; *P* = 1).

In addition to the above results from the planned comparisons that motivated our study, an unexpected finding also emerged from our multi-species analysis. Specifically, we observed significant convergent amino acid enrichment between the echolocating bats and both the African elephant (*Loxodonta africana*) and the mouse (*Mus musculus*) to the exclusion of the non-echolocating bats (**Figure 3; Supplemental Table 1**). For example, African elephants shared 31 out of 7,540 amino acids with *Rhinolophus* to the exclusion of *Eidolon*, compared to only 8 amino acids shared with *Eidolon* to the exclusion of *Rhinolophus*; the resulting convergence ratio is significantly greater than expected by chance (convergence ratio = 3.9; Fisher’s Exact test, *P* = 0.000289, Bonferroni-corrected *P* = 0.013). Likewise, mice shared 32 out of 8,196 amino acids with *Rhinolophus* to the exclusion of *Eidolon* yet only 4 with *Eidolon* to the exclusion of *Rhinolophus* (convergence ratio = 8; *P* = 1.9E-6; Bonferroni-corrected *P* = 8.3E-5).

## Discussion

We did not identify any signatures of genomic convergence between aye-ayes and true echolocators at the seven genes related to auditory processing that we examined. While our analysis cannot account for the potential presence of non-identical amino acid changes that are nonetheless convergent in biochemical function, the finding of strong convergent evolution for specific amino acid changes at these loci between echolocating dolphins and bats but not between aye-ayes and either of these lineages suggests that aye-aye tap-foraging auditory adaptations represent functionally differentiated evolutionary innovations compared to echolocating bats and dolphins.

However, a surprising result from our analysis suggests that the auditory gene convergence phenomenon may not be unique to echolocating bats and dolphins. Specifically, we observed unexpectedly significant enrichments for convergent amino acids between echolocating bats and mouse and elephant. Interestingly, mice have been observed to communicate in the ultrasound during courtship and mating (Musolf et al. 2010), with a frequency comparable to that used by many echolocating bats (~70 kHz; White et al. 1998), and outside the range of aye-aye auditory ability (Ramsier and Dominy 2012). Perhaps the observed convergent evolution between mouse and echolocating bats at these loci is spurred by the demands of high-ultrasound processing rather than echolocation behavior? However, if this were correct, then convergence might also be expected between mouse and dolphins, who also have high-ultrasound processing demands via echolocation, yet such a result was not observed (**Figure 3**). In contrast to mice, African elephants are known for their ability to hear *low* frequency sounds in the infrasonic range (<20 Hz; Payne et al. 1986; Langbauer et al. 1991) rather than the high frequency ultrasound (1-200 kHz) of echolocating bats (Au and Simmons 2007). Thus, perhaps the apparent convergent evolution between echolocating bats and both elephants and mice at these loci indicates that some amino acid changes at these genes could play a more general role in auditory signal conduction at frequency extremes, perhaps with some unknown, similar underlying biology among these taxa, rather than only at high frequencies as part of echolocation behavior.

## Acknowledgments

This research was supported by funds from the Penn State College of the Liberal Arts and the Huck Institutes of the Life Sciences (to GHP), and the Pennsylvania State University Graduate Fellowship (to RB). The Duke Lemur Center kindly provided the diademed sifaka DNA samples (this is DLC publication # XXXXX), and Parker *et al.* (2013) provided access to the gene alignment data from their paper. We thank Craig Praul, Candace Price, and the Penn State Huck Institutes of the Life Sciences Genomics Core for generating the Illumina sequencing data, Stephen Johnson, Kate Thompson, David Villalta, Richard George, Alexis Sullivan, Arslan Zaidi, James Fisher, Stephen King, and Kristen Alldrege for discussions and feedback on the study and manuscript, and Emily Davenport for suggestions on the GitHub repository. Computational resource instrumentation was funded by the National Science Foundation through Grant OCI–0821527.

